# Exploratory noise governs both flexibility and spontaneous errors and is regulated by cocaine

**DOI:** 10.1101/328872

**Authors:** R. Becket Ebitz, Brianna J. Sleezer, Hank P. Jedema, Charles W. Bradberry, Benjamin Y. Hayden

**Author notes:** Corresponding author and lead contact: Becket Ebitz, Department of Neuroscience, University of Minnesota, Minneapolis MN 55455.

## Abstract

In many cognitive processes, lapses (spontaneous errors) are attributed to nuisance processes like sensorimotor noise or disengagement. However, some lapses could also be caused by exploratory noise: behavioral randomness that facilitates learning in changing environments. If so, strategic processes would need only up-regulate (rather than generate) exploration to adapt to a changing environment. This view predicts that lapse rates should be correlated with flexibility because they share a common cause. We report that when macaques performed a set-shifting task, lapse rates were negatively correlated with perseverative error frequency. Furthermore, chronic exposure to cocaine, which impairs cognitive flexibility, increased perseverative errors, but, surprisingly, improved overall performance by reducing lapse rates. We reconcile these results with a model in which cocaine decreased exploration by deepening attractor basins corresponding to rules. These results support the idea that exploratory noise contributes to lapses, meaning that it affects rule-based decision-making even when it has no strategic value.

## INTRODUCTION

Decision-makers can implement arbitrary rules (i.e. stimulus-response mappings) and flexibly change them when contingencies change (Miller and Cohen, 2001; Wallis et al., 2001). Yet even sophisticated decision-makers occasionally fail to implement well-learned rules. Why do these lapses occur? In general, lapses of rule adherence, are tacitly dismissed as the result of ancillary nuisance processes, such as memory deficits, sensorimotor noise, or disengagement (McVay and Kane, 2009; Reason, 1990; Van der Linden et al., 2003; Weissman et al., 2006). An alternative view is that some lapses occur because of the same adaptive processes that allow rule learning and cognitive flexibility in a changing environment. Determining whether lapse rates are somehow linked to the capacity for flexibility could provide insight into psychiatric illnesses in which lapse rates are abnormal (e.g. (Ciesielski and Harris, 1997; Floresco et al., 2009; Heinrichs and Zakzanis, 1998)), and into the basic mechanisms of rule use.

In changing environments, decision-makers mostly exploit valuable strategies, but they also occasionally explore, i.e. deviate from valuable strategies to sample alternatives (Berg and Brown, 1972; Ebitz et al., 2018; Kaelbling et al., 1996; Pearson et al., 2009; Sutton and Barto, 1998; Wilson et al., 2014). In many algorithms for exploration, the likelihood of exploration depends on uncertainty or the value of exploring (Daw et al., 2006; Kaelbling et al., 1996; Sutton and Barto, 1998). In these *phasic* algorithms, exploration occurs most often when reducing perseveration has the greatest benefit. In *tonic* algorithms, conversely, the decision does not depend on uncertainty or the value of exploration (Kaelbling et al., 1996; Sutton and Barto, 1998). Although tonic exploration may appear suboptimal, it eliminates the need to calculate the value of exploration at every time step, is robust to errors in calculating the value of exploration, and it can perform nearly as well as phasic exploration in many circumstances (Dayan and Daw, 2008; Ebitz et al., 2018; Sutton and Barto, 1998). Tonic exploration also has costs: when the environment is stable it produces errors of rule adherence that have no immediate strategic benefit. That is, it produces lapses.

It remains unclear whether exploration occurs even when it has no strategic value. One way to address this question is by looking at behavior in a “change-point” task (Behrens et al., 2007; Nassar et al., 2012; O’Reilly et al., 2013; Wilson et al., 2010). Change-point tasks have stable periods—in which there is no uncertainty and exploratory noise has no strategic benefit— and also rapid changes in reward contingencies that require adaptation and learning. If exploration occurs tonically—if it does contribute to lapses—then spontaneous lapses during stable periods should be related to the ability to discard a rule. That is, across animals and days, lapse rates should be negatively correlated with perseverative errors. An alternative hypothesis is that exploration is phasic, generated only at change points. If so, then lapse rates would not be correlated with perseverative errors (because they are caused by different processes), or perhaps positively correlated (because they are both errors).

Furthermore, if lapse rates and adaptation at change points are both caused by tonic exploration, then it should be possible to identify an intervention that simultaneously alters both behaviors because it regulates this underlying common cause. One candidate intervention is chronic cocaine exposure, which reduces cognitive flexibility (Bechara, 2005; Everitt and Robbins, 2005; Jentsch et al., 2002; Lucantonio et al., 2012; Robbins and Everitt, 1999). Cocaine abusers make more perseverative errors in classic set-shifting tasks such as the Wisconsin Card Sort Task (WCST; (Beatty et al., 1995; Colzato et al., 2009; van der Plas et al., 2009; Woicik et al., 2011)). Both rodents and monkeys exposed to cocaine show deficits in reversal learning (Porter et al., 2011; Schoenbaum et al., 2004) and fail to change behavior in the face of aversive outcomes (Vanderschuren and Everitt, 2004). This inflexibility may contribute to the cycle of abuse in cocaine users (Everitt and Robbins, 2005; Robbins and Everitt, 1999; Turner et al., 2009).

If cocaine exposure regulates tonic exploration, then it should not only cause perseverative errors, but also decrease lapse rates. It should simultaneously decrease flexibility yet improve performance in set-shifting tasks. Indeed, at least one observational study reported that human cocaine abusers performed better in the WCST, compared to controls (Hoff et al., 1996). However, it remains unclear whether chronic cocaine is sufficient to simultaneously reduce lapse rates and increase perseverative errors. Addressing this question has the potential to reconcile seemingly paradoxical results in the cocaine literature, and, at the same time, to address a fundamental question about whether lapses are caused by the same tonic exploration process that facilitates adaptation and learning.

Therefore, we examined behavior of rhesus macaques performing the cognitive set shifting task (CSST) (Moore et al., 2005; Sleezer and Hayden, 2016; Sleezer et al., 2016, 2017; Yoo et al., 2018), a primate analogue of the WCST, both before and after exposure to cocaine. This task is ideal to address the present question because it combines a change point task with a rule-based decision-making task. Consistent with tonic exploration, we found evidence of a common cause of lapse rates during stable periods and flexibility following change points. Moreover, cocaine not only reduced flexibility, but simultaneously and proportionally decreased lapse rates, suggesting that cocaine regulates tonic exploration. Finally, we fit a model to the dynamics of behavior, in which cocaine decreased exploration via deepening the attractor basins that correspond to rule states. Together these results suggest that exploration occurs tonically and may be well-described as variation in the depth of attractor basins corresponding to rule states.

## RESULTS

Two macaques performed 147 sessions of a primate analogue of the WCST (the CSST (Moore et al., 2005; Sleezer and Hayden, 2016; Sleezer et al., 2016, 2017; Yoo et al., 2018); **Figure 1A**) before and after chronic self-administration of cocaine (n = 89 baseline sessions before cocaine administration, monkey B: n = 62, monkey C: n = 27; n = 58 post-cocaine sessions after, monkey B: 33, monkey C: 25). In a trial, monkeys were sequentially offered three choice options that differed in both color and shape (drawn from nine possible combinations of three colors and three shapes). On each trial, one of the six stimulus features was associated with reward. The rewarded rule was chosen randomly and remained fixed until a rule change was triggered (by successful completion of 15 trials). Rule changes were not cued. We have not previously examined this data in the way presented below nor have we previously reported the results of cocaine exposure.

**Figure 1.**
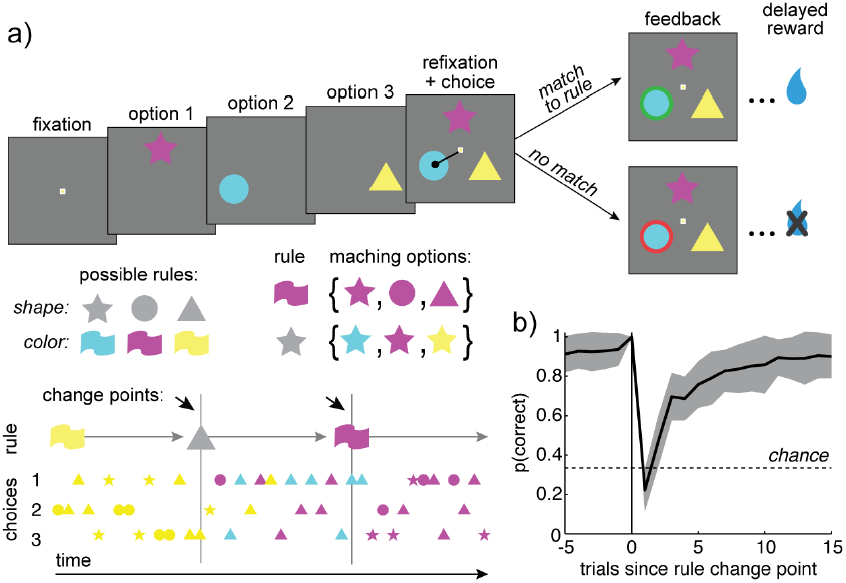
Task design and baseline behavior. A) The CCST task. Three options, which differed in both shape and color were sequentially presented. Choosing an option that matched the rewarded rule produced a green outline around the chosen option and a reward. Choosing either of the other two options produced a red outline and no reward. Middle row, left: Rules could be any of the three shapes or any of the three colors. Right: The options that matched a rule were the set of stimuli that shared the rule’s feature. Bottom: After the monkeys achieved 15 correct choices, the rewarded rule changed, which forced the monkeys to search for the new rule. B) Percent correct as a function of trials before and after rule changes. The 0^th^ trial is the last trial before the rule changed. Gray shading +/- STD.

Monkeys chose the most rewarding option frequently (81.4% of trials ± 6.5% STD across sessions, monkey B = 83.9% ± 5.8% STD, monkey C = 77.1% ± 5.7% STD; average of 576 trials per session, 470 rewarded) and adapted quickly to rule changes (**Figure 1B**). Most errors were perseverative (repeated either the color or shape of the previous option; 64 ± 8.5% STD across sessions; average of). Pre-cocaine sessions were collected after 3 months of training. We observed no measurable trend in performance across the pre-cocaine sessions (**Figure 2A**; percent correct, GLM with terms for main effects of monkey and session number, session number beta = 0.0002, p = 0.6, df = 86, n = 89). Thus, performance had reached stable levels before data collection began.

**Figure 2:**
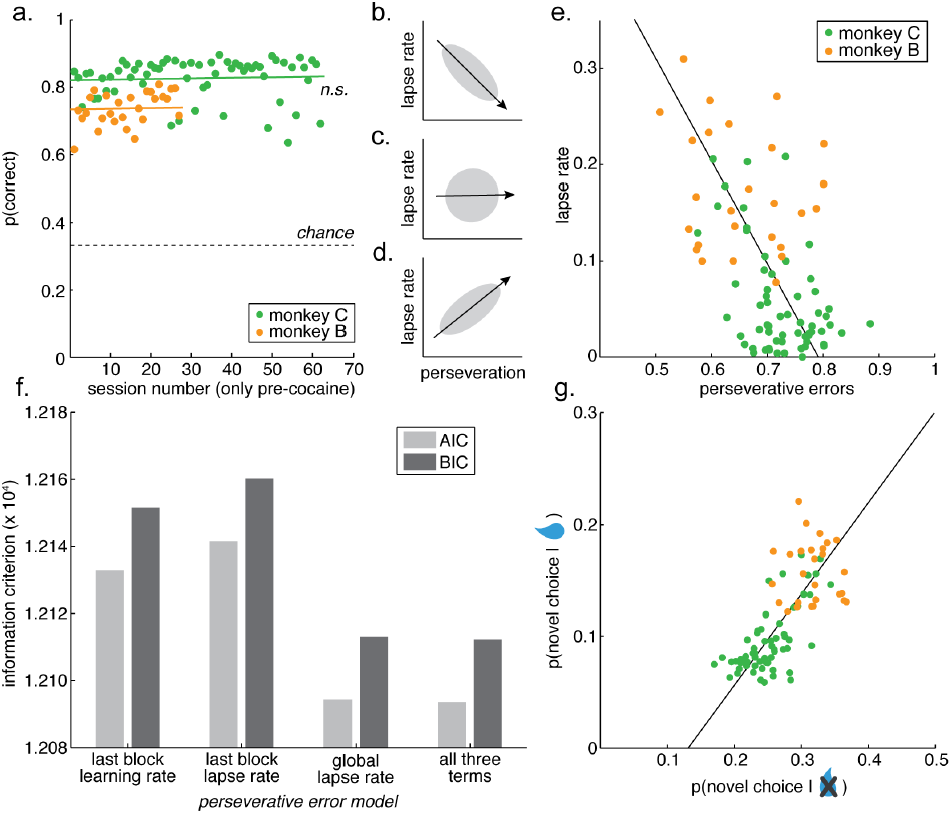
Behavior in baseline sessions. A) Percent correct as a function of session-number in the baseline sessions, plotted separately for monkey C (green dots) and monkey B (orange). Lines are GLM fits for each monkey (Results). n.s. = not significant. B-D) Cartoon depicting the possible relationships between lapse rates and perseverative errors under different hypotheses. B) Some spontaneous lapses are caused by the same process that facilitates learning and reduces perseveration at change points. C) Lapses and perseveration are caused by different underlying error processes. D) Lapses and perseveration are both caused by a common error process, such as disengagement. E) The observed relationship between lapses in the 10 trials proceeding change points and perseverative errors in the 5 trials after change points. F) Model comparison to determine whether perseverative errors are more closely related to the rate of learning or lapse rate in the last block or to the global lapse rate in that session. G) The correlation between the likelihood of novel choices (matching neither the last color nor last shape), given reward delivery and omission. Best fit lines = ordinary least squares.

### Relationship between lapse rates and perseverative errors

Lapses are a failure to adhere to a good policy when the environment has not changed. Perseverative errors are the continued adherence to a bad policy when the environment has changed. These two behaviors could be related (or unrelated) for a variety of reasons.

We considered three hypotheses, each of which predicted a different relationship between lapses during stable periods and perseverative errors after change points. *First*, if spontaneous errors of rule adherence (lapses) are caused by the same process that helps to discard a rule when it is no longer rewarded (e.g. tonic exploratory noise) then lapse rates would be negatively correlated with perseverative errors across sessions (**Figure 2B**). *Second*, if lapses and perseverative errors are regulated by different processes (e.g. if lapses occur because of a transient memory deficit, while perseverative errors occur because of a failure of inhibitory control), then the frequency of lapses and perseverative errors would not be correlated (**Figure 2C**). *Third*, if some nuisance process causes both types of errors (e.g. disengagement or fatigue), then lapses and perseverative errors would be positively correlated (**Figure 2D**).

We compared perseverative errors in the five trials after change points (when learning was maximal; **Figure 1B**) with lapse rates in the ten trials before change points (a non-overlapping subset of trials in which learning had reached asymptote). Lapse rates and perseverative errors were negatively correlated (**Figure 2E**; both monkeys: Pearson’s r = −0.52, p < 0.0001, n = 89). This was not a trivial consequence of a performance offset between the monkeys: the effect was strongly significant just within the monkey in which we had more baseline data (monkey C: n = 62 sessions, r = −0.45, p < 0.0002; same sign in monkey B: n = 27 sessions, r = −0.26, p = 0.25). A negative correlation between lapses and perseverative errors indicates that the rate of lapses in rule adherence is positively correlated with the ability to discard a rule when it is no longer rewarded.

Lapse rates in one epoch cannot directly cause flexibility in another epoch (or vice versa), so this correlation implies that both behaviors share some common, underlying cause. One possibility is tonic exploration, which would cause monkeys to occasionally sample an alternative to the current best option, regardless of change points. Another possibility is a failure to learn, which would cause lapses (because the rule is never discovered) and reduce perseverative errors (because a rule that is never discovered is cannot persevere). The failure-to-learn view predicts that perseverative errors in one block should be best explained by the lapses in the immediately preceding block. However, the probability of perseverative errors in each individual block was best explained by the global lapse rate for the session, not to the lapse rate or the rate of learning in the previous block (**Figure 2F**; see Methods; last-block lapse rate model: log likelihood = −6063.4, AIC = 12133, BIC = 12152; last-block learning rate model: log likelihood = −6067.8, AIC = 12142, BIC = 12160; global lapse rate model: log likelihood = −6044.2, AIC = 12094, BIC = 12113; best model = global lapse rate model, all other AIC and BIC weights < 0.0001). Thus, the negative correlation between lapse rates and perseverative errors was not due to a failure to learn in some blocks, but instead to some global common cause, such as tonic exploration.

In this task, the outcome of the previous trial provides perfect information about whether or not that choice was correct. If monkeys were rewarded on the last trial, then either the color or shape of the last choice matched the rewarded rule and the best response is to repeat either the color or shape or both in the next trial. Conversely, if the monkeys were not rewarded, then neither the color or shape of the last choice was consistent with the rewarded rule and the best response is to choose a novel option—one that matches neither the color nor the shape of the previous choice. However, tonic exploration would sometimes cause monkeys to choose novel options following reward delivery—when it is clearly incorrect to do so. Indeed, the monkeys did choose novel options after both reward delivery (monkey B: 15.8% novel choices, monkey C: 9.6%) and omission (monkey B: 31.6% novel choices, monkey C: 25.2%). However, tonic exploration not only predicts that these choices should occur, but that their frequency should be governed by a common underlying process. That is, the frequency of novel choices after reward delivery should be correlated with the frequency of novel choices after reward omission. Indeed, these choices were strongly correlated (**Figure 2G**; Pearson’s r = 0.72, p < 0.0001, n = 89). This was individually significant within the animal in which we had more baseline sessions (monkey C: n = 62 sessions, r = 0.68, p < 0.0001; monkey B: n = 27 sessions, r = −0.04, p = 0.9). Thus, the monkeys’ decisions to deviate from choice history—to try something new—also co-varied, regardless of whether or not that was correct, consistent with a common cause.

### Cocaine self-administration

The variability in the baseline behavior suggested a common process regulating the decision to deviate from a rule, regardless of whether or not it is correct to do so. If this is true, then it should be possible to co-regulate lapses and perseverative errors by regulating this process. Therefore, we next allowed both monkeys to self-administer cocaine—exposure to which is known to affect the ability to adapt to a changing environment (Bechara, 2005; Everitt and Robbins, 2005; Jentsch et al., 2002; Lucantonio et al., 2012; Porter et al., 2011; Robbins and Everitt, 1999).

Monkeys self-administered cocaine through an implanted venous port (see Methods). Briefly, for 3 hours each day, 5 days a week, over a total of 6 to 7 weeks (monkey B: 50 days, monkey C: 42 days), monkeys were placed in front of a touch screen display and pressed a centrally located cue a set number of times (see Methods), which resulted in cocaine infusion. Monkeys initially underwent self-administration training (10 days). During this time, the cumulative dose of cocaine self-administered per day increased from 0.8 mg/kg to 4 mg/kg at 3 responses/reward (FR3), followed by a ramp-up period to 30 responses/reward (FR30; 7 days at 4 mg/kg), after which we began examining behavioral data during chronic cocaine exposure. We collected behavior in the morning, while monkeys self-administered cocaine in the afternoon in a separate session (with a minimum of 1 hour of home cage time in between). This experimental design allowed us to determine the long-term effects of chronic cocaine self-administration without the drug “on board” at the time of testing. Over all self-administration sessions, monkey B administered a cumulative total of 179.9 mg/kg of cocaine, while monkey C administered 153.2 mg/kg cocaine.

### Effects of cocaine on behavior

Because chronic cocaine exposure is associated with decreased flexibility and increased perseveration, we first asked whether cocaine administration changed the proportion of perseverative errors. It did (**Figure 3A**; fraction of all errors that were perseverative, post cocaine compared to pre, t-test: p < 0.0001, t(145) = 6.13, mean increase in fraction perseverative errors = 7.7%, 95% CI = 5.1% to 10.0%; monkey B: p < 0.0001, t(58) = 7.70; monkey C: p < 0.0001, t(85) = 6.99). One concern in any study of chronic drug use is that practice alone could change behavior and appear to be a drug effect. To test for this possibility, we developed a generalized linear model (GLM) to differentiate between the effects of drugs and practice (see Methods). There was no effect of practice on perseverative errors (β_2_ = 0.003, p = 0.7) and including a term for session number did not change the magnitude of the effect of cocaine (β_1_ = 0.097, p < 0.0001), indicating that practice explained little, if any, change in perseverative errors in post-cocaine sessions.

**Figure 3:**
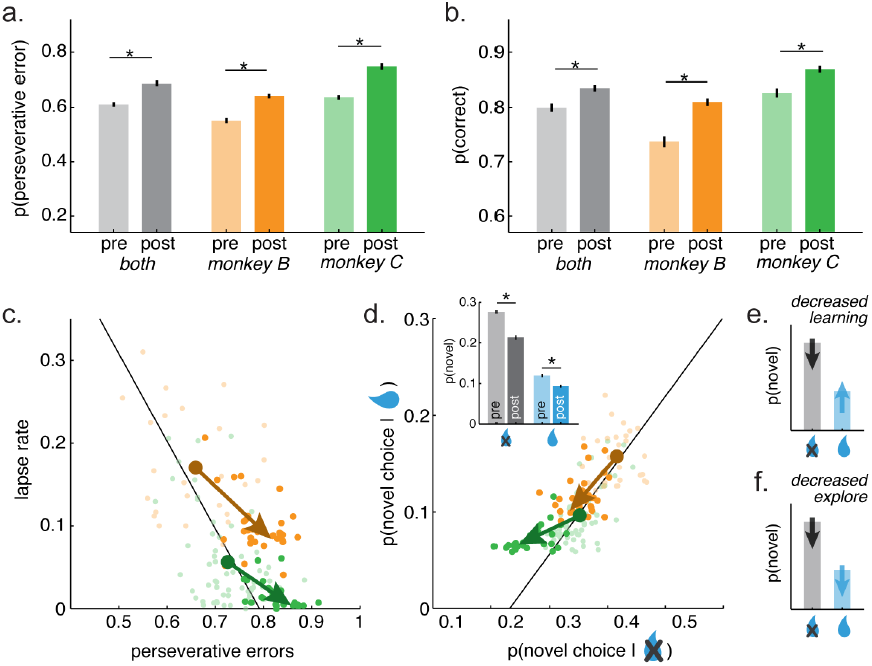
Changes in CSST behavior after cocaine administration. A) The probability of perseverative errors before and after cocaine treatment (before = light, after = dark), plotted together for both monkeys (gray) as well as separately for monkey B (orange bars) and monkey C (green). Error bars +/- SEM throughout and ^*^ p < 0.05, two-sample t-test. B) Same as A, for the percent of total correct trials in the pre- and post-cocaine sessions. C) Cocaine’s effects on the relationship between spontaneous lapses and perseverative errors. Same as 2E, but now illustrating post-cocaine sessions (dark) and pre-cocaine sessions (light). The vectors reflect the shift in the mean with cocaine for monkey B (orange) and monkey C (green). D) Cocaine’s effects on the relationship between novel choices after reward delivery (ordinate) and omission (abscissa). Same as 2G, but with the conventions of 3C. Inset) Change in novel choice probability, plotted separately for reward omission (gray) and delivery (blue). Pre-cocaine = light, post cocaine = dark. E) An illustration of the hypothesis that cocaine decreases learning rates. We would have expected to see a decrease in the difference between novel choices following reward delivery and reward omission in D, inset. F) Same as E, for the hypothesis that cocaine decreases exploration, in which case it would reduce all novel choices, without regard to previous reward outcome.

If cocaine increased perseveration by decreasing tonic exploration, then it might also improve overall performance in this set-shifting task by reducing lapse rates. Cocaine reduced whole-session error rates (**Figure 3B**; percent correct, post cocaine compared to pre, t-test: p < 0.001, t(145) = 3.36, mean increase = 3.6%, 95% CI = 1.5% to 5.7%; monkey B: p < 0.0001, t(58) = 6.30; monkey C: p < 0.002, t(85) = 3.22). Again, session number did not affect accuracy (β_2_ = 0.001, p = 0.9) and accounting for session number only increased the apparent magnitude of the effect of cocaine (compare 3.6% change to β_1_ = 0.054, p < 0.0005). This was likely driven by the substantial decrease in the frequency of lapses in the 10 trials before change points (figure 3C; two-sample t-test; monkey B: p < 0.0001, t(58) = 5.57, mean difference = 7.1%, 95% CI = 4.6% to 9.7%; monkey C: p < 0.0006, t(85) = 3.59, mean = 4.0%, 95% CI = 1.8% to 6.2%).

The hypothesis that cocaine regulates a common cause of flexibility and lapses makes a strong prediction: that cocaine should simultaneously shift lapses and perseverative errors along the axis on which they endogenously co-vary (line in Figure 2E). This is because this axis reflects the consequences of any common cause on both lapses and perseverative errors. Therefore, any modulation of this common cause should be constrained to shifts along this manifold. Therefore, we measured the projection of the pre- and post-cocaine sessions onto the line along which the two behaviors endogenously co-varied (see Methods). Cocaine significantly shifted behavior along this axis (two-sample t-test, both monkeys: p < 0.0001, t(145) = 7.60, mean shift in standardized projection = 0.77, 95% CI = 0.57 to 0.98). The effect was significant and of comparable magnitude in both monkeys (monkey B: p < 0.0002, t(58) = 4.09, mean = 0.72, 95% CI = 0.37 to 1.07; monkey C: p < 0.0001, t(85) = 5.48, mean = 0.68, 95% CI = 0.44 to 0.93). This is precisely the effect that we would expect if cocaine regulated the underlying cause of both behaviors.

Next, we asked whether cocaine had similar effects on monkeys’ decisions to deviate from their own previous policy. That is, the probability of novel choices (**Figure 2G**). A decrease in tonic exploration would decrease the likelihood of novel choices regardless of previous reward outcome, so asked whether chronic cocaine decreased novel choices following both reward delivery and omission. Cocaine decreased the probability of novel choices both after reward omission (when novel choices were the best option, **Figure 3D**; two-sample t-test, both monkeys, p < 0.0001, t(145) = 6.16, mean change = −5.1%, 95% CI = −3.4 to −6.7%; monkey B: p < 0.0001, t(58) = 7.99; monkey C: p < 0.0001, t(85) = 8.57; not due to practice β_1_ = −0.057, p< 0.0001; β_2_ = −0.008, p = 0.1) and after reward delivery (when novel choices were the worst option, both monkeys, p < 0.006, t(145) = 2.83, mean change = −1.7%, 95% CI = −0.5 to −2.9%; monkey B: p < 0.0001, t(58) = 6.97; monkey C: p < 0.001, t(85) = 3.50; not due to practice β_1_ = −0.024, p < 0.002; β_2_ = −0.005, p = 0.2). It is important to note that if cocaine decreased learning (i.e. the effect of reward on behavior), then it would decrease the difference between choices following reward delivery and reward omission (**Figure 3E**). However, cocaine instead decreased the probability of novel choices, regardless of reward outcome, consistent with tonic exploration (**Figure 3F**).

If these effects are due to cocaine’s effects on tonic exploration, then cocaine should simultaneously alter the probability of novel choices regardless of previous outcome. That is, cocaine should shift novel choice probability along the axis of endogenous co-variability between rewarded and non-rewarded trials (line in Figure 2G). It did so (Figure 3D: two-sample t-test, both monkeys, p < 0.0001, t(145) = 5.78, mean change = 0.49, 95% CI =0.32 to 0.66; monkey B: p < 0.09, t(58) = 1.73; monkey C: p < 0.0001, t(85) = 7.85). Thus, cocaine appeared to regulate the probability of making novel choices directly, rather than modulating the effect of rewards on novel choices. Because tonic exploration would produce novel choices both when they are useful and when they are not, this result is consistent with the idea that chronic cocaine down-regulates tonic exploration.

### Hidden Markov model

We previously developed a method to identify whether individual choices are exploratory or exploitative based on a hidden Markov model (HMM) (Ebitz et al., 2018). Here, we extend this model to dissociate exploratory choices from choices that were made while using rules (**Figure 4A**). We chose this framework for two reasons. First, because HMMs are useful for interring the latent “states” that underlie a sequence of observations (such as the explore and rule goal states that underlie the sequences of choices here). Second, because HMMs describe behavior in terms of the dynamics of these underlying states, which allowed us to analyze how cocaine changed the dynamics of explore and rule goal states.

**Figure 4:**
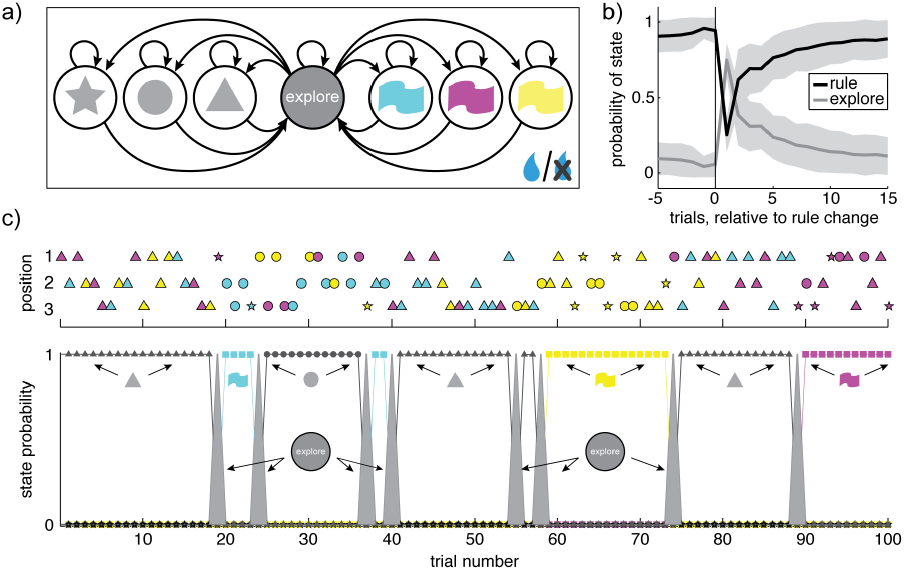
Hidden Markov model (HMM) design and fit to behavior. A) The structure of the HMM, with one latent state for each possible rule, plus one latent “explore state”. Emissions (not shown) match the rule in the rule states, and are randomly allocated during the explore state. The box around the model indicates that this model has multiple “plates”, which depend on the reward of the previous trial (bottom right). That is, each path (transition probability between states) depends on whether the animal was or was not rewarded on the previous trial. B) The posterior probability of explore states and any of the rule states (1-p(search)) is illustrated as a function of trials relative to change points in the rewarded rule. Shading: +/- STD. C) Top: A sequence of 300 chosen options, separated vertically by whether the chosen option was in location 1, 2, or 3. Bottom, the state probabilities from a fitted HMM. Colored lines with colored boxes correspond to the color-rule states (blue, yellow, and magenta). Black lines with black shape icons correspond to shape-rule states (triangle, circle, square). The gray shaded line corresponds to the explore state probability.

We reasoned that rule-states would only generate choices that matched the rule, but while exploring monkeys would choose many different kinds of choices. Therefore, we next asked whether there was evidence of these different dynamics in behavior. Indeed, there were distinct dynamics associated with repeated choices within a feature dimension (i.e. following a rule) and rapid samples across feature dimensions (i.e. exploring; **Figure S1**). These rapid samples occurred more frequently than expected, suggesting a distinct exploratory state (**Figure S2)**. We also found that the duration of choice runs depended on reward (**Figure S3**). To account for this, we extended model so the outcome of the last trial affected the probability of transitioning between states (“transmissions”, see Methods; (Bengio and Frasconi, 1995)). The final HMM (see **Methods**) qualitatively reproduced the reward-dependent state durations (**Figure S3**) and the latent states inferred by this model successfully differentiated choices that occurred due to each of these dynamics (example in **Figure 4B**). In addition, the latent states inferred by the model were strongly aligned with the change points in the task, indicating that the model was most likely to identify choices as exploratory at precisely the time when the monkeys were actually searching for a new rule (compare **Figure 4C** and **Figure 1B**).

Next we asked whether the model was capable of reproducing the major behavioral effects of cocaine. We fit one model to all the baseline sessions and a second model to the post-cocaine sessions, then simulated observations from each model. The changes in model parameters across the baseline and post-cocaine sessions were sufficient to reproduce the major behavioral results: an increase in both task performance (**Figure 5A**; mean increase in percent correct = 14.5%, 95% CI = 12.8 to 16.1%, p < 0.0001, t(145) = 17.70) and perseverative errors (**Figure 5B**; mean increase in percent perseverative errors = 4.8%, 95% CI = 3.9 to 5.8%, p < 0.0001, t(145) = 9.89). Thus, the model captured the main effects of cocaine on behavior.

**Figure 5:**
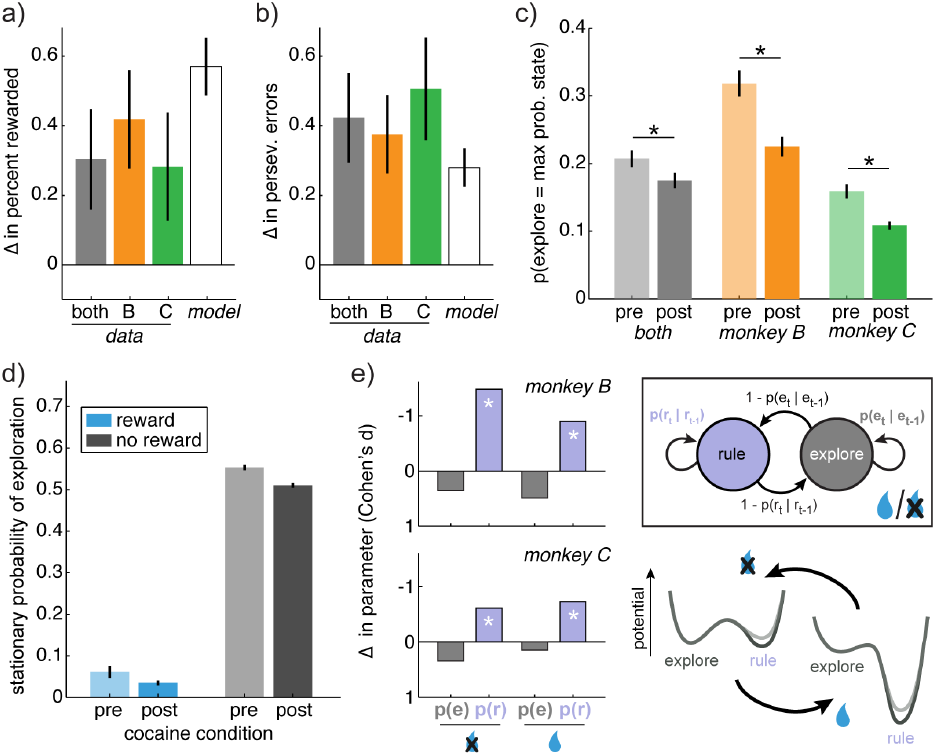
HMM predictions and effects of cocaine on model behavior. A) The increase in the probability correct after cocaine. Plotted separately for both monkeys together (gray bar), monkey B (orange) and monkey C (green), next to the increase in probability correct in simulated data from the model (white bar). Bars: Satterthwaite approximation of the +/ 99 CI. B) Same as A, for change in perseverative errors. C) The probability that exploration was identified as the most probable cause of each choice, before and after cocaine. Gray=both monkeys together, orange=monkey B, green=monkey C. Bars +/- SEM. D) The stationary probability of the explore state, given the outcome of the previous trial (rewarded=blue, not rewarded=gray) and the cocaine condition (pre=before cocaine, post=after). E) Effect of cocaine on the the 4 free parameters in the model (top left). Change in parameters (Cohen’s d, post-cocaine minus baseline) in monkey B (top) and monkey C (bottom). ^*^ p < 0.05, t-test (see Table 1). Note that the slight decrease in the probability of staying in exploration was likely due to practice (see Results). Bottom right) A cartoon illustrating the effect of cocaine on model parameters (see Table 1) in terms of an attractor landscape. Here, exploration and rule adherence correspond to some local minima in a behavioral landscape, across which the monkeys move stochastically. Reward outcomes act to shift the baseline landscape (light line) from strongly favoring rule adherence following reward delivery (left) to a slight preference for exploration following reward omission (right; compare to panel D). Cocaine (dark line) globally increases the duration of rule-states, which suggests that it specifically deepens the attractor basin corresponding to rules, regardless of reward outcome.

### Cocaine reduces HMM-inferred exploration

Next, we asked whether cocaine affected the probability of exploration, as inferred from the model using a standard algorithm (Viterbi algorithm). One model was fit to each session, then each choice was labeled by its max a posteriori latent state. The monkeys had different levels of exploration, but within each monkey, there were fewer explore-state choices in post-cocaine treatment sessions, compared to baseline sessions (**Figure 5C**; monkey B: p < 0.0002, t(58) = 4.03, mean change = −9.3%, 95% CI = −4.7 to −13.9%; monkey C: p < 0.004, t(85) = 3.01, mean = −5.0%, 95% CI = −1.7 to −8.4%; not due to practice: β_1_ = 0.052, p < 0.03; β_2_ = 0.011, p = 0.3). Thus, monkeys explored less often after cocaine delivery, consistent with the idea that cocaine alters tonic exploration.

### Effects of cocaine on model dynamics

The stationary distribution of a HMM is the equilibrium probability distribution over states (Murphy, 2012). Here, the HMM’s stationary distribution is the relative occupancy of explore-states and rule-states that we would expect after infinite realizations, given the outcome of the last trial (see Methods). That is, it provides a measure of the energetic landscape of the behavior the model is fit to. If a state has very low potential energy—if its basin of attraction is deep—then we will be more likely to observe the process in this state, and the stationary distribution will be shifted towards this state (Ambegaokar, 2017). Therefore, we will refer to the stationary distribution probability of exploration as the “relative depth” of exploration.

As expected, reward delivery reduced the relative depth of explore states (**Figure 5D;** and increased the relative depth of the rule states; see Methods; β_1_ = −0.49, p < 0.0002). Cocaine also decreased the relative depth of explore states (β_2_ = −0.05, p < 0.02). There was a significant offset between monkeys (β_4_ = −0.05, p < 0.0002) and no effect of practice (β_5_ = 0.0003, p = 0.4) or interaction between reward and cocaine (β_3_ = 0.016, p = 0.4). This suggested that cocaine uniformly altered the depth of exploration, rather than the effect of reward on exploration. To test this, we asked whether the effect of cocaine on explore state depth differed after reward delivery, compared to reward omission. There was no significant difference after controlling for the expected effect of differing baselines (see Methods; paired t-test: p = 0.9, t(144) = −0.09, mean change = 1%, 95% CI = −25% to 23%). Moreover, the depth of exploration was correlated across reward outcome within the baseline sessions (both monkeys: r = 0.38, p < 0.0001, n = 89) and cocaine delivery did not disrupt these correlations (both monkeys: Pearson’s r = 0.23, p < 0.005, n = 147). Thus, cocaine uniformly decreased the relative depth of exploration, regardless of reward outcomes.

### Effects of cocaine on model parameters

Did cocaine reduce the relative depth of explore states by increasing the absolute depth of exploration or by increasing the absolute depth of rule states? To arbitrate between these interpretations, we next asked how cocaine changed the parameters of the model. The model had 4 parameters (**Figure 5E**), reflecting the probability of staying in each of the two states (explore and the generic rule state) following the two outcomes (reward delivery and omission). If cocaine largely affected the probability of staying in exploration, then that would suggest that cocaine specifically decreased the depth of explore states. This is because the average dwell time in a state (that is, the inverse of the rate of leaving that state) has a natural relationship to the energetic depth of that state, relative to the energy barrier between states (Hänggi et al., 1990). Alternatively, if cocaine largely affected the probability of staying in a rule, then that would suggest that cocaine specifically increased the depth of rule states. We also considered a third possibility: that cocaine had different effects following reward delivery and omission—i.e. decreasing the depth of rules after reward omission, but increasing depth of exploring after reward delivery. This last effect would be hard to reconcile with the idea of a unified effect on tonic exploration.

Within each monkey, there were significant changes in the same two model parameters in post-cocaine sessions (**Table 1**). Cocaine increased the probability of staying in rule states following reward omission (monkey B: p < 0.0001, t(58) = 5.69; monkey C: p < 0.02, t(85) = 2.57; not due to practice: β_1_ = 0.070, p < 0.04, β_2_ = 0.027, p = 0.1) and cocaine increased the probability of staying in rule states following reward delivery (monkey B: p < 0.001, t(58) = 3.45; monkey C: p < 0.003, t(85) = 3.06; not due to practice: β_1_ = 0.004, p < 0.01, β_2_ = 0.0002, p=0.8). Cocaine had no significant effect on the depth of explore states following either reward omission (β_1_ = −0.004, p > 0.9) or reward delivery (β_1_ = 0.03, p = 0.7). However, there was a trend towards a decrease in the depth of explore states with practice in both conditions (omission: β_2_ = −0.03, p = 0.1, delivery: β_2_ = −0.06, p = 0.09). Thus, the weight of evidence suggests that cocaine selectively deepened rule states (Figure 5E): it decreased tonic exploration via increasing the tendency to adhere to a rule, regardless of reward outcomes.

**Table 1:** Effects of cocaine on model parameters. Mean parameter estimate (standard deviation) across all models. p(e_t_) = probability of exploration. p(r_t_) = probability of rule. Bold: significant change in post-cocaine sessions, relative to baseline within each monkey: ^*^ p < 0.05, ^**^ p < 0.005, ^***^ p < 0.0001, t-test (see Results for test statistics).

| Parameter |  | Monkey B |  | Monkey C |  |
| --- | --- | --- | --- | --- | --- |
|  |  | Baseline | Post-cocaine | Baseline | Post-cocaine |
| Reward | $p(r_t\vert r_{t-1})$ | 0.978 (0.008) | **0.984 (0.006)\*\*** | 0.995 (0.005) | **0.998 (0.002)\*\*** |
| | $p(e_t\vert e_{t-1})$ | 0.73 (0.17) | 0.64 (0.21) | 0.30 (0.30) | 0.25 (0.25) |
| No reward | $p(r_t\vert r_{t-1})$ | 0.02 (0.07) | **0.19 (0.14)\*\*\*** | 0.04 (0.11) | **0.11 (0.12)\*** |
| | $p(e_t\vert e_{t-1})$ | 0.28 (0.16) | 0.22 (0.17) | 0.18 (0.14) | 0.14 (0.12) |

## DISCUSSION

We found that spontaneous lapses and perseverative errors were not independent observations, but instead were inversely related across monkeys and sessions. This was not a trivial consequence of the monkeys’ ability to learn the rewarded rule. Instead, there was a global common cause of both lapses and perseverative errors, which meant that the two types of error inversely co-varied along a one-dimensional manifold. Moreover, chronic cocaine—a perturbation known to decrease flexibility and increase perseveration (Bechara, 2005; Everitt and Robbins, 2005; Jentsch et al., 2002; Lucantonio et al., 2012; Porter et al., 2011; Robbins and Everitt, 1999)—did not uniquely increase perseverative errors, but instead shifted the animals along this manifold. That is, cocaine produced a concomittant decrease in lapse rates. To understand these results, we fit and analyzed a HMM, which revealed that cocaine decreased exploration via deepening attractor basins corresponding to rule states.

These results suggest that the same process that facilitates flexibility in a dynamic environment is responsible for at least some spontaneous lapses in rule adherence when the environment is stable. That is, these results suggest that exploratory noise is tonically present, and causes deviations from established decison policies, both when these deviations are useful and when they are not.

### Relationship to previous theories of lapses and flexibility

We are not proposing that tonic exploratory noise is categorically different from other processes that are typically implicated in lapses, such as disengagement, memory deficits, sensorimotor noise, or attentional or executive disengagement (McVay and Kane, 2009; Reason, 1990; Van der Linden et al., 2003; Weissman et al., 2006). Instead, we propose that these may be valid psychological descriptions of the effect that exploratory noise has on behavior.

What, then, is exploratory noise in the brain? Exploratory decisions are associated with sudden disruption in the choice-predictive organization of populations of neurons the prefrontal cortex (Ebitz et al., 2018). It is possible that this disorganization reflects a disruption of the prefrontal attractor dynamics that are thought to underpin working memory (Brody et al., 2003; Chaudhuri and Fiete, 2016; Compte et al., 2000; Kopec et al., 2015; Wimmer et al., 2014), motor control (Li et al., 2016), decision-making (Machens et al., 2005; Wang, 2002, 2008), and executive control (Ardid and Wang, 2013; Rougier et al., 2005). These dynamics could allow these regions to influence the behavior of lower-order circuitry (Ebitz and Moore, 2017), perhaps via amplifying the information available to the prefrontal cortex (Wang, 2008). Disrupting these dynamics, then, could have a range of psychological effects, which might be unified if thought of as randomizing behavior with respect to information or policies held in the prefrontal cortex.

On the surface, the link between lapses and perseverative errors that we report here may appear to conflict with previous views of errors in similar tasks as reflecting separate and dissociable cognitive processes. Many modern theories of flexibility view perseveration as measuring the (in)ability to inhibit a previous rule and lapses as measuring the (in)ability to either maintain a rule or to inhibit distraction from irrelevant options (Barceló, 1999; Barceló and Knight, 2002; Block et al., 2007; Floresco et al., 2006, 2009; Ragozzino, 2007). The present results can be reconciled with these theories if increasing depth of a rule makes it both easier to maintain and harder to inhibit. Increasing the depth of a rule could also decrease distraction, either by regulating the frequency of exploration or by regulating the strength of rule processes that otherwise outcompete distraction. There is precedent for the view that internal states linked to exploration (Jepma and Nieuwenhuis, 2011) also predict increased distraction (Ebitz and Platt, 2015; Mather and Sutherland, 2011). Moreover, tonic exploration almost certainly cannot explain all errors of task performance and it remains likely that increases in the number of lapses following other perturbations arise from changes in other cognitive processes (Barceló, 1999; Barceló and Knight, 2002; Block et al., 2007; Floresco et al., 2006, 2009; Ragozzino, 2007).

### Relationship to previous views of cocaine

The fact that cocaine administration increases perseverative responding is well-established (Bechara, 2005; Everitt and Robbins, 2005; Jentsch et al., 2002; Lucantonio et al., 2012; Porter et al., 2011; Robbins and Everitt, 1999). However, here cocaine simultaneously improved overall performance in a set-shifting task—the exact type of task in which perseveration should interfere with performance. At least one previous study reported that chronic cocaine use correlates with improved performance in a set shifting task (Hoff et al., 1996). Here, we replicate both results within the same animals in a causal study. We also reconcile both results with a simple formalism—a hidden Markov model in which cocaine deepened the attractor basins corresponding to rule states. Together, these results suggest that cocaine acts to stabilize rules, making it harder to break out from using a rule, either spontaneously or in response to feedback from the environment.

The perseverative effects of chronic cocaine use have previously been interpreted as a shift from goal-directed, action-outcome or model-based control systems to habitual, stimulus-response or model-free control systems (Bechara, 2005; Everitt and Robbins, 2005; Jentsch and Taylor, 1999; Jentsch et al., 2002; LeBlanc et al., 2013; Lucantonio et al., 2012; Robbins and Everitt, 1999; Robinson and Berridge, 1993). The present results support these views. In particular, these results support the influential hypothesis that cocaine shifts monkeys into a model-free decision-making regime, in which learning is slow and choices are habitual (Lucantonio et al., 2012). Although cocaine had no effect on the animals’ sensitivity to rewards (there was no change in the difference in behavior following reward omission and delivery), it did increases the *hysteresis* of response policies—that is, the tendency to persist in a policy simply because you have been using it (Lau and Glimcher, 2005). This is consistent with previous observations that cocaine selectively interferes with learning when a previously-learned response must be overcome (Jentsch et al., 2002; Lucantonio et al., 2012; Porter et al., 2011) and observations that cocaine directly increases the probability of repeating responses (LeBlanc et al., 2013; Stout et al., 2004). We are not the first to note the link between exploratory noise and the balance between model-free and model-based decision-making (Dayan and Daw, 2008) and the present results suggest that regulating tonic exploratory noise may be the mechanism by which cocaine causes a shift towards model-free decision-making.

### Basic insights into the mechanistic bases of flexibility

The lawful relationship between lapses and perseverative errors was not an artificial consequence of cocaine exposure. Instead, cocaine shifted behavior along the axis of endogneous co-variability that already existed between these error types: tonic exploration was a meaningful parameter that was controlled by cocaine administration, not introduced by it. Thus, the neurobiological targets of cocaine exposure may be promising targets for understanding the neural basis of tonic exploration.

One important cortical target of chronic cocaine administration is the orbitofrontal cortex (OFC) (Lucantonio et al., 2012; Schoenbaum et al., 2004; Stalnaker et al., 2009): a region that is implicated in rule encoding (Baeg et al., 2009; Sleezer et al., 2016; Tsujimoto et al., 2011; Wallis et al., 2001; Yamada et al., 2010). Orbitofrontal damage leads to a deficit in maintaining performance during stable, steady periods in the WCST (Stuss et al., 2000) and results in choice behavior that is consistent with an inability to learn or maintain rules (Walton et al., 2010). Of course, other cortical regions are also likely to contribute to regulating flexibility, particularly the anterior cingulate cortex (Ebitz and Hayden, 2016; Ebitz and Platt, 2015), and there are functional and structural difference in both the cingulate and the OFC in chronic cocaine exposure (Baeg et al., 2009; Franklin et al., 2002). Thus, these region are an important target for future studies of both cognitive flexibility and the effects of drugs of abuse.

Cocaine exposure also has profound effects on the brains’ neuromodulatory landscape. Chronic cocaine alters the dopamine (DA) (Bradberry et al., 2000; Burchett and Bannon, 1997; Gifford and Johnson, 1992; Hurd et al., 1990; Pettit et al., 1990), norepineprine (NE) (Beveridge et al., 2005; Burchett and Bannon, 1997; Macey et al., 2003), acetylcholine (ACh) (Gifford and Johnson, 1992; Hurd et al., 1990), and serotonin (Burchett and Bannon, 1997) systems. ACh, DA and NE, in particular, have been previously implicated in regulating exploratory decision-making (Aston-Jones and Cohen, 2005; Doya, 2002; Yu and Dayan, 2005). Moreover, lesions of ACh interneurons in the dorsomedial striatum may be sufficient to produce a change in lapse rates and perseverative errors simular to those reported here (Aoki et al., 2015). The effects of cocaine here support hypotheses linking these neuromodulatory systems to exploration, but the hypothesis that cocaine regulates exploration via regulating these neuromodulatory systems will need to be tested empirically.

### Conclusions

Why would exploratory noise influence behavior even when it has no strategic benefit? One possibility is that tonic exploration may have conferred such substantial benefits over evolutionary time that our brains evolved to maintain it even when it has no value in the moment. What benefits might these be? For one, up-regulating an existing stochastic noise process may simply be a more efficient use of metabolic resources than overcoming an embedded strategy de novo. For another, tonic exploratory noise could reduce the energetic and/or computational costs of deciding *when* to explore. In tonic exploration there is no need to calculate the value of exploration at each time step (Dayan and Daw, 2008).

Oddly, tonic exploration could also facilitate rule adherence by eliminating this calculation. In artificial intelligence literature, temporally-extended behavioral policies—known as “options”—can speed planning, reduce computational costs, and increase the capacity for complex and abstract goals (Sutton et al., 1999). Clearly there are parallels between options and cognitive rules (Miller and Cohen, 2001). It is notoriously difficult, however, for agents to learn to use options because it is always more valuable to re-evaluate the choice of option at each time step than to commit to one (Harb et al., 2017; Sutton et al., 1999). This is because commitment to an option imposes opportunity costs, even when the value of the alternatives is very low (Harb et al., 2017; Lloyd and Dayan, 2018). Tonic exploration would solve this problem because it ensures that alternatives to the current policy are occasionally sampled, but without the need to calculate the value of alternatives or indeed the need to represent the opportunity cost of extended commitment. Moreover, allowing agents to only probabilistically commit to a rule lowers the opportunity cost of commitment (Lloyd and Dayan, 2018). Thus, tonic exploratory noise may be an important part of how we evolved the ability to apply rules, as well as an intrinsic part of how we apply rules today.

## Methods

### General surgical procedures

All animal procedures were approved by the University Committee on Animal Resources at the University of Rochester and were conducted in accordance with the Public Health Service’s Guide for the Care and Use of Animals. Two male rhesus macaques (Macaca mulatta) served as subjects. The animals had previously been implanted with small prosthetics for holding the head (Christ Instruments), which allowed us to monitor eye position and use this as the response modality. These procedures have been described previously (Strait et al., 2014). To allow for chronic cocaine self-administration, we also implanted a subcutaneous vascular access port (VAP) in these animals (Access Technologies, Skokie, IL, USA), which was connected via an internal catheter to the femoral vein. Additional details of the VAP implantation procedure have been reported previously (Bradberry et al., 2000; Wojnicki et al., 1994). The VAP allowed monkeys to self-administer cocaine daily, and obviated the need for chemical or physical restraint, which might have unintended consequences for behavior. Animals received appropriate analgesics and antibiotics after all procedures, per direction of University of Rochester veterinarians. The animals were habituated to laboratory conditions and trained to perform oculomotor tasks for liquid reward before training on the conceptual set shifting task (CCST) began. Both animals participated in laboratory tasks for at least two years before the present experiment.

### Self-administration protocol

The monkeys sat in a primate chair placed in a behavioral chamber with a touchscreen (ELO Touch Systems, Menlo Park, CA, USA). Syringe Pump Pro software (Version 1.6, Gawler, South Australia) controlled and monitored a syringe pump (Cole Parmer, Vernon Hills, IL, USA), which delivered cocaine into the monkeys’ VAP. Monkeys pressed a centrally located visual cue on the touchscreen to obtain venous cocaine injections (cocaine provided by National Institutes of Drug Abuse, Bethesda, MD, USA), delivered in a 5 mg/ml solution at a rate of 0.15 ml/s. Monkeys were acclimated to cocaine self-administration across ten days of training, during which the response requirement and dose increased from 3 responses/reward (FR3) and 0.1 mg/kg (0.8 mg/kg of cocaine daily) to 30 responses/reward (FR30) and 0.5 mg/kg (4 mg/kg of cocaine daily). Monkeys were given 3 hours to complete infusions each day (in practice, monkeys typically completed the all 8 infusions within 1-2 hours). Monkeys self-administered cocaine 5 days a week.

### Behavioral task

Specific details of this task have been reported previously (Sleezer and Hayden, 2016; Sleezer et al., 2016, 2017; Yoo et al., 2018). Briefly, the present task was a version of the CSST: an analogue of the WCST that was developed for use in nonhuman primates (Moore et al., 2005). Task stimuli are similar to those used in the human WCST, with two dimensions (color and shape) and six specific rules (three shapes: circle, star, and triangle; three colors: cyan, magenta, and yellow; figure 1A). Choosing a stimulus that matches the currently rewarded rule (i.e. any blue shape when the rule is blue; any color of star when the rule is star) results visual feedback indicating that the choice is correct (a green outline around the chosen stimulus) and, after a 500 ms delay, a juice reward. Choosing a stimulus that does not match the current rule results in visual feedback indicating that the choice is incorrect (a red outline), and no reward is delivered after the 500 ms delay.

The rewarded rule was fixed for each block of trials. At the start of each block, the rewarded rule was drawn randomly. Blocks lasted until monkeys achieved 15 correct responses that matched the current rule. This meant that blocks lasted for a variable number of total trials (average = 22.5), determined by both how long it took monkeys to discover the correct objective rule and how effectively monkeys exploited the correct rule, once discovered. Block changes were uncued, although reward-omission for a previously rewarded option provided noiseless information that the reward contingencies had changed.

On each trial, three stimuli were presented asynchronously, with each stimulus presented at the top, bottom left, or bottom right of the screen. The color, shape, position, and order of stimuli were randomized. Stimuli were presented for 400 msec and were followed by a 600-msec blank period. (The blank period was omitted from Figure 1A because of space constraints). Monkeys were free to look at the stimuli as they appeared, and, though they were not required to do so, they typically did (Sleezer and Hayden, 2016). After the third stimulus presentation and blank period, all three stimuli reappeared simultaneously with an equidistant central fixation spot. When they were ready to make a decision, monkeys were required to fixate on the central spot for 100 msec and then indicate their choice by shifting gaze to one stimulus and maintaining fixation on it for 250 msec. If the monkeys broke fixation within 250 milliseconds, they could either again fixate the same option or could change their mind and choose a different option (although they seldom did so). Thus, the task allowed the monkeys ample time to deliberate over their options, come to a choice, and even change their mind, without penalty of error.

### General data analysis techniques

Data were analyzed with custom MATLAB scripts and functions. All t-tests were two-sample, two-sided tests, unless otherwise noted. All generalized linear models (GLMs) included a dummy-coded term to account for a main effect of monkey identity (1 for monkey B, 0 for monkey C) and were fit to session-averages, rather than individual trials. One session (1/147) was excluded from these analyses because one of its transmission matrices did not admit a stationary distribution. No data points were excluded for any other reason. Observation counts for each analysis are reported in figure legends and/or Results.

### Differentiating the effects of cocaine treatment from practice

Task performance reached stable levels in both monkeys before the baseline, pre-cocaine sessions began (figure 2A). Nevertheless, we were concerned that putative effects of cocaine self-administration might instead be trivial consequences of the increased experience with the task in the post-cocaine sessions. Any effect of cocaine treatment would produce a step change in behavior that was aligned to the start of cocaine administration. Conversely, the effects of practice would change gradually across sessions. Therefore, to determine whether individual behavioral effects were due to practice or cocaine, we fit the following GLM to the session-averaged behaviors of interest:

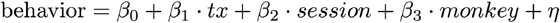

Where “tx” is a logical vector indicating whether the session was conducted before or after chronic cocaine self-administration (a step change term) and “session” was a vector of session number within the experiment for each monkey (a gradual ramping term). One additional term “monkey” accounted for the random effect of monkey identity, and the model included the standard intercept and noise terms (β_0_ and η, respectively). Thus, β_1_ captured any offset due to chronic cocaine administration, while β_2_ captured any effect of practice for each analysis.

### Probability of novel choices

Only 3 of the 9 possible stimuli (i.e. 9 combinations of 3 colors and 3 shapes) were available on each trial, so the likelihood of repeating choices that shared neither feature was constrained by the available options. Therefore, we calculated the monkeys’ probability of choosing each number of feature repeats as the total number of times a certain number of features was repeated, divided by how many times it was possible to repeat that number of features. Both terms were calculated within session.

### Hidden Markov Model

In the HMM framework, choices (y) are “emissions” that are generated by an unobserved decision process that is in some latent, hidden state (z). Latent states are defined by both the probability of each emission, given that the process is in that state, and by the probability of transitioning to or from each state to every other state. Straightforward extensions of this framework allow inputs, such as rewards, to influence state transitions (Bengio and Frasconi, 1995), in which case the latent states can be thought of as a kind of discretized value function.

The observation model for each hidden state is the probability choosing each option when the process that state. These emissions models differed across the two broad classes of states in the model—the explore states and rule states—based on the fact that there were two different dynamics in the choice behavior: one reflecting random choosing while exploring and one reflecting long staying durations due to persistent rules (Figures S1 and S2). Therefore, the observation model for any choice option *n* during explore states was:

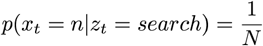

Where N is the number of stimuli that were presented (i.e. N=3). During rules, the observation model was conditioned on a match between each stimulus and the current rule:

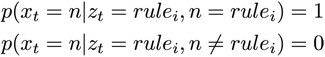

The latent states in this model are Markovian meaning that they are time-independent. They depend only on the most recent state (z_t_) and most recent reward outcome (u_t_):

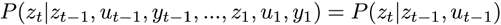

This means that the probabilities of each state transition are described by reward-dependent transmission matrix, A_k_ = {a_i,j_}_k_ = P(z_t_ = j | z_t-1_ = i, u_t-1_ = k) where k ∈{rewarded, not rewarded}. There were 7 possible states (6 rule states and 1 explore state) but parameters were tied across rule states such that each rule state had the same probability of beginning (from exploring) and of sustaining itself. Similarly, transitions out of explore were tied across rules, meaning that it was equally likely to start using any of the 6 rules after exploring. Because monkeys could not divine the new rule following a change point and instead had to explore to discover it, transitions between different rule states were not permitted. The model assumed that monkeys had to pass through explore in order to start using a new rule, even if only for a single trial. Thus, each plate k of the transition matrix had only two parameters, meaning there were a total of 4 parameters in the reward-dependent model.

The model was fit via expectation-maximization using the Baum Welch algorithm (Bilmes, 1998; Murphy, 2012). This algorithm finds a (possibly local) maxima of the complete-data likelihood, which is based on the joint probability of the hidden state sequence Z and the sequence of observed choices Y, given the observed rewards U:

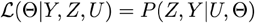

The complete set of parameters Θ includes the observation and transmission models, discussed already, as well as an initial distribution over states, typically denoted as π. Because monkeys had no knowledge of the correct rule at the first trial of the session, we assumed the monkeys began in the explore state. The algorithm was reinitialized with random seeds 100 times, and the model that maximized the observed (incomplete) data log likelihood was ultimately taken as the best for each session. The model was fit to individual sessions, except to generate simulated data, in which case one model was fit to all baseline sessions and a second to all post-cocaine sessions. To decode latent states from choices, we used the Viterbi algorithm to discover the most probable a posteriori sequence of latent states (Murphy, 2012).

To simulate data from the model, we created an environment that matched the monkeys’ task (choices between 3 options with 2 non-overlapping features and a randomly selected rewarded rule that changed after 15 correct trials). We then probabilistically drew latent states and choice emissions as the model interacted with the environment. The only modification to the model for simulation was that the choice of rule state following a explore state was constrained to match one of the two features of the last choice, chosen at randomly.

### Stationary distribution

To gain insight into how cocaine changed the likelihood of rule states following reward delivery and omission, we examined the stationary distributions of the model. The transmission matrix of a HMM is a system of stochastic equations describing probabilistic transitions between each state. That is, each entry of a transmission matrix reflects the probability that the monkeys would move from one state (e.g. exploring) to another (e.g. using a rule) at each moment in time. In this HMM, there were two transmission matrices, one describing the dynamics after reward delivery and one describing the dynamics after reward omission. Moreover, because the parameters for all the rule states were tied, each transition matrix effectively had two states—an explore state and a generic rule-state that described the dynamics of all rule states. Each of these transition matrices (A_k_) describes how the entire system—an entire probability distribution over explore and rule states—would evolve from time point to time point given the outcome of the previous trial, k. You can observe how these dynamics would change any probability distribution over states π by applying the dynamics to this distribution:

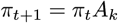

Over many iterations of these dynamics, ergodic systems will reach a point where the state distributions are unchanged by continued application of the transmission matrix as the distribution of states reaches its equilibrium. That is, in these systems, there exists a stationary distribution, π^*^, such that:

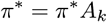

If it exists, this distribution is a (normalized) left eigenvector of the transition matrix A_k_ with an eigenvalue of 1, so we solved for this eigenvector to determine the stationary distribution of each A_k_, if it had one. (Only one of the A_k_ matrices did not admit a stationary distribution, so this session was not included in analyses related to this measure.)

### Analyzing stationary distributions

To determine how cocaine affected the relative depth of exploration and the generic rule state, we constructed a GLM. The model included terms to describe the effects of reward, cocaine, and the interaction between the two on the depth of exploration. This interaction allowed the model to describe a phasic, reward-dependent effect of cocaine on the depth of exploration, if it were present:

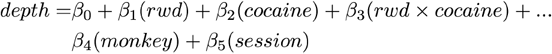

The model thus accounted for any offset between monkeys (“monkey”, 1 for monkey B, 0 for monkey C) or practice effects (“session”). It also included terms to describe the effects of reward (“rwd”, 1 for reward delivery, 0 for omission), cocaine (“cocaine”, 1 for pre-cocaine baseline sessions, 0 for post-cocaine sessions), and the interaction between reward and cocaine. This allowed the model to describe a phasic, reward-dependent effect of cocaine on model dynamics or a tonic, reward-independent form of exploration.

### Comparing changes in probabilities

We calculated log odds ratios to compare the magnitude of changes in probability when baseline probabilities differed. Because probabilities are bounded, they are necessarily nonlinear transformations of an unbounded latent process of interest. This means that a fixed change in an underlying linear process can produce very different magnitude changes in probability, depending on the baselines. For intuition, picture a logistic function—a typical nonlinear transformation used to covert linear observations into probabilities. The effect of an equivalent change in the x-axis on the y-axis is depends on the baseline position on the x-axis: an identical shift on the x-axis has a large effect on y when x starts close to the midpoint of the function, but a small effect on y when x starts close to either end. The logit transformation linearizes the relationship between different observed probabilities because it is the inverse of the the logistic function:

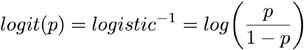

The difference between log odds (also known as the log odds ratio) then provides us with a linearized measure of effect magnitude (less sensitive to differing baseline levels). It is:

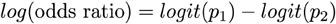

## Acknowledgements

The authors would like to thank Nicola Grissom and Habiba Azab for comments on the manuscript, Daniel Takahashi for invaluable discussion, Marc Mancarella, Meghan Pesce, and Giuliana Loconte for technical help and assistance with animal care and husbandry. Support provided by the National Institute on Drug Abuse (R01-DA038106) and the Brain & Behavior Research Foundation (NARSAD award to BYH).

## Author Contributions

BJS and BYH designed the behavioral experiment; BJS, BYH, HPJ and CWB designed the cocaine protocol; HPJ and BJS performed surgeries; BJS collected the data with guidance from BYH, HPJ, and CWB; BJS, BYH, and RBE formulated the hypotheses; RBE analyzed the data; BYH secured funding; RBE drafted the manuscript, which all authors edited.

## Supplemental Figures and References

**Supplemental Figure 1.**
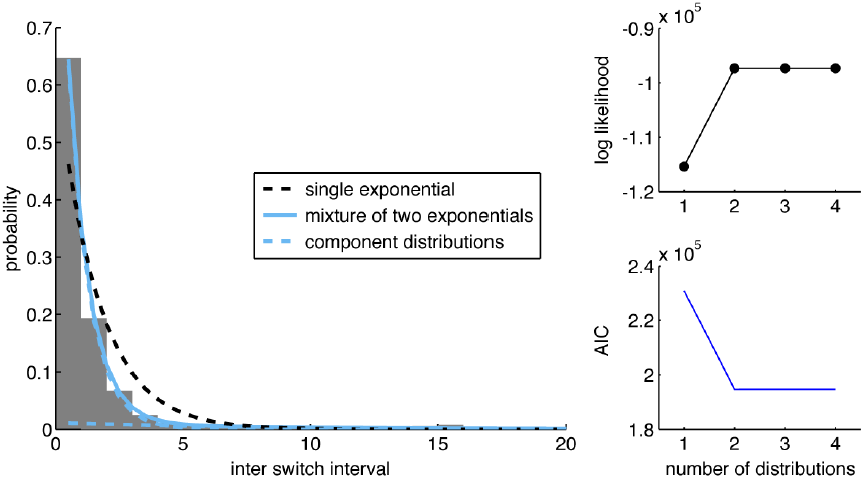
Hidden Markov Model development (related to figures 4 and 5). To determine whether an HMM was an appropriate descriptive model for this dataset, we first asked whether there were different behavioral dynamics that might correspond to using a rule and exploring. One way to do this is to examine the distribution of runs of repeated choices within some choice dimension (Ebitz, Albarran, & Moore, 2018). If monkeys are exploiting a rule, then they would have to repeatedly choose options that are consistent with this rule. During a rule, runs of repeated choices—or interswitch intervals—would be long. However, exploration, monkeys need to briefly sample the options to determine whether or not they are currently rewarded. That is, during exploration runs of repeated choices should be very brief: on the order of single trials. To the extent that choice runs end because of stochastic events (an assumption of the HMM framework), inter-switch intervals will be exponentially distributed (Berg, 1993). Moreover, if there are multiple latent regimes (such as exploring and rule-following), then we would expect to see inter-switch intervals distributed as a mixture of exponential distributions, because choice runs have a different probability of terminating in each latent regime. The distribution of inter-switch intervals (n interswitch intervals = 49,059) resembled an exponential (**left**), but was better described by a mixture of two discrete exponential distributions (blue lines; 1 exponential: 1 parameter, log-likelihood = −142077.0, AIC = 284156.1, AIC weight < 0.0001, BIC = 284165.6, BIC weight < 0.0001; (Burnham and Anderson, 2003)) than a single distribution (black line; 2 exponential: 3 parameters, log-likelihood = −119773.2, AIC = 239552.4, AIC weight = 1, BIC = 239580.7, BIC weight = 1). Adding additional exponential distributions did not improve model fit (r**ight**), suggesting that there were only two regimes (3 exponentials: 5 parameters, log-likelihood = −119773.2, AIC = 239556.4, AIC weight < 0.14, BIC = 239603.7, BIC weight < 0.0001; 4 exponentials: 7 parameters, log-likelihood = −119773.2, AIC = 239560.4, AIC weight < 0.02, BIC = 239626.6, BIC weight < 0.0001). The best-fitting model, the two-exponential mixture had one long-latency component (half life = 9.0), consistent with a persistent rule-following response mode. It also had one short latency component (half life 1.4; consistent with random choice between 3 options).

**Supplemental Figure 2.**
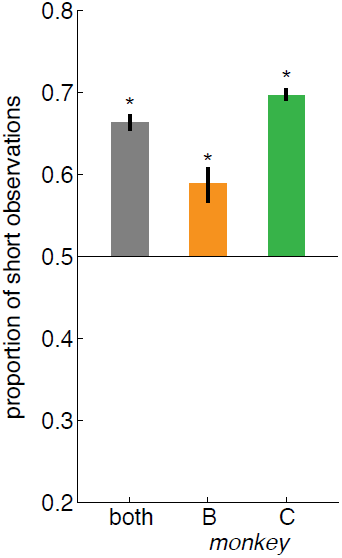
Short choice runs occur more frequently than expected (related to figures 4 and 5). Because rules only operated on either the color or shape of the option, we quantified the duration of inter-switch intervals independently within the color and shape domains (i.e. a magenta star choice followed by a magenta circle choice be counted as part of the same choice run in the color domain, but would part of different choice runs in the shape domains). This meant that choices would inevitably be randomized within one feature domain during repeated choices in the other domain. Thus, the existence of a mode with a short half-life is not sufficient evidence of short-latency search dynamics. However, if randomization in the other domain was the sole cause of short duration samples, then observations from the short sampling mode would occur exactly as frequently as observations from the persistent mode. However, short choice runs occurred more frequently than expected. To determine this, we calculated the expected time in each state as the product of the average run length in that state and the probability of being in that state. Then, we normalized the expected time in the short state by the sum of expected times in all states. That is, this measure would be at 0.5 if observations from the short state were equally as frequent, and greater than 0.5 if they were more frequent. The expected number of short state observations was significantly greater than 0.5 (both subjects, paired t-test, p < 0.0001, t(88) = 17.02; subject B: p < 0.0003, t(26) = 4.18; subject C, p < 0.0001, t(61) = 27.6), indicating that both subjects had more frequent short duration samples than would be expected if those short duration samples were merely caused by choices along a different dimension. Thus, both subjects exhibited strong evidence for a separate search state, in which they made short duration runs of choices to the different options.

**Supplementaly Figure 3.**
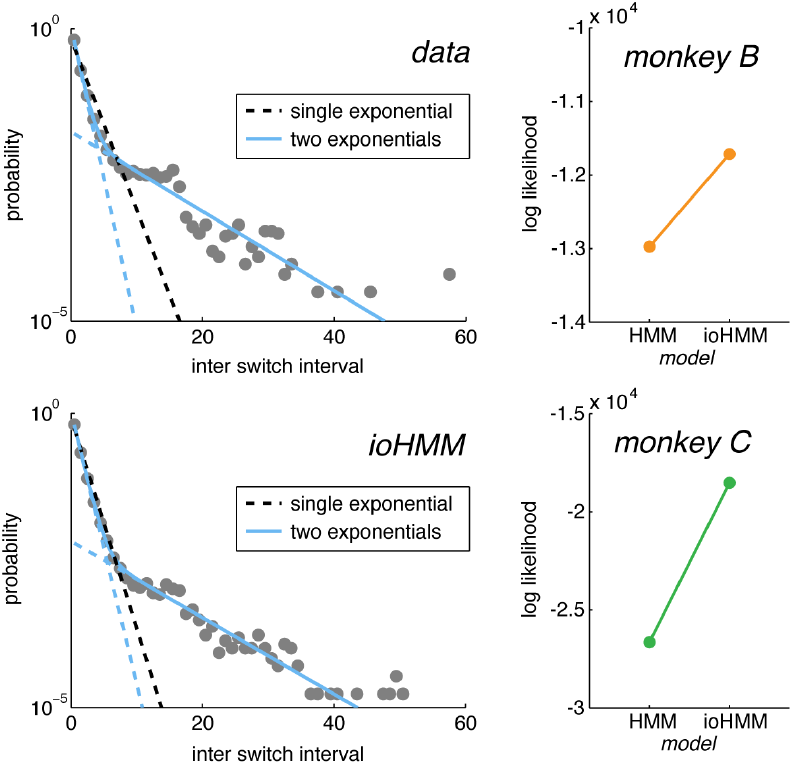
An input-output HMM accounts for reward-dependent decisions (related to figures 4 and 5). Inter-switch intervals were largely exponential—consistent with the Markovian assumptions of an HMM—and we observed different search and rule dynamics. However, it is important to note that in the log plot (**top left**), there were significant deviations from the predictions of simple exponential mixture model. These were likely due to the changes in reward contingencies that were triggered each time 15 correct trials were completed. To account for the obvious dependence on reward, we extended a simple 2 parameter HMM model to allow state transition probabilities to depend on previous reward outcomes (Bengio and Frasconi, 1995). Accounting for this reward dependence (4-parameter ioHMM) qualitatively reproduced these dynamics (**bottom left**) and quantitatively improved model fit in both monkeys (**right**; both monkeys: 2 parameter HMM, log-likelihood = −39614, 4 parameter ioHMM, log-likelihood = −30240, log-likelihood ratio test: statistic 18749, p < 0.0001; monkey B: HMM, log-likelihood = −12973, ioHMM = −11714, log-likelihood ratio test: statistic = 2518.7 p < 0.0001; monkey C: HMM, log-likelihood = −26641, ioHMM = −18526, log-likelihood ratio test: statistic = 16230, p < 0.0001).

